# FilterDCA: interpretable supervised contact prediction using inter-domain coevolution

**DOI:** 10.1101/2019.12.24.887877

**Authors:** Maureen Muscat, Giancarlo Croce, Edoardo Sarti, Martin Weigt

## Abstract

Predicting three-dimensional protein structure and assembling protein complexes using sequence information belongs to the most prominent tasks in computational biology. Recently substantial progress has been obtained in the case of single proteins using a combination of unsupervised coevolutionary sequence analysis with structurally supervised deep learning. While reaching impressive accuracies in predicting residue-residue contacts, deep learning has a number of disadvantages. The need for large structural training sets limits the applicability to multi-protein complexes; and their deep architecture makes the interpretability of the convolutional neural networks intrinsically hard. Here we introduce FilterDCA, a simpler supervised predictor for inter-domain and inter-protein contacts. It is based on the fact that contact maps of proteins show typical contact patterns, which results from secondary structure and are reflected by patterns in coevolutionary analysis. We explicitly integrate averaged contacts patterns with coevolutionary scores derived by Direct Coupling Analysis, reaching results comparable to more complex deep-learning approaches, while remaining fully transparent and interpretable. The FilterDCA code is available at http://gitlab.lcqb.upmc.fr/muscat/FilterDCA.

**Author summary:** The *de novo* prediction of tertiary and quaternary protein structures has recently seen important advances, by combining unsupervised, purely sequence-based coevolutionary analyses with structure-based supervision using deep learning for contact-map prediction. While showing impressive performance, deep-learning methods require large training sets and pose severe obstacles for their interpretability. Here we construct a simple, transparent and therefore fully interpretable inter-domain contact predictor, which uses the results of coevolutionary Direct Coupling Analysis in combination with explicitly constructed filters reflecting typical contact patterns in a training set of known protein structures, and which improves the accuracy of predicted contacts significantly. Our approach thereby sheds light on the question how contact information is encoded in coevolutionary signals.

## Introduction

The prediction of protein structure using amino-acid sequence information has seen important progress in the last decade. This progress is substantially based on two subsequent methodological advances: unsupervised sequence-based coevolutionary models, and subsequent structure-based supervision using deep learning.

First, global coevolutionary methods like Direct Coupling Analysis (DCA) [1, 2], PSICOV [3] or Gremlin [4, 5] have allowed to identify direct coevolutionary couplings by modeling the sequence variability found in sufficiently large homologous protein families. It was found that the largest coevolutionary couplings correspond, with high probability, to residue-residue contacts inside or between proteins [6, 7]. The potential for tertiary and quaternary protein structure prediction is evident, and many interesting cases have been published [8–11]. However, the fully unsupervised character of coevolutionary models is limiting their wide-scale applicability: they are based only on statistical models of sequence ensembles, and do not make use of available structural information. This fact leads, in turn, to a restricted number of predicted contacts, in particular in the case of protein families containing only a limited number of sequences.

In the case of tertiary structure, this problem has been solved more recently. Benefitting from the large numbers of proteins with experimentally determined structures available in the PDB, and from the large amount of residue-residue contacts (and non-contacts) in each structure, supervised machine-learning techniques have allowed to substantially increase the accuracy in predicting residue-residue contacts and even distances. In particular convolutional neural networks (CNN) based on deep learning have shown success, cf. the performance of methods like RaptorX [12], DeepMetaPSICOV [13] and AlphaFold [14] in the last editions of the CASP structure prediction competition [15].

Despite their impressive predictive performance, CNN and other deep neural architectures have some important disadvantages. First, they need to be trained on large datasets. While these are available for tertiary protein structures, the applicability to inter-protein contact prediction and protein-complex assembly has remained limited, even if promising results have been found when training was performed on intra-protein contacts, but testing on inter-protein contacts [16]. Second, deep learning is intrinsically hard to interpret. It remains normally unclear how deep networks extract information from data, and how they use it to annotate residue pairs as contacts or non-contacts.

The architecture of CNN may give a hint. Instead of looking individually to residue pairs, their first layer applies convolutional filters to a neighbourhood of each residue pair. Protein contact maps are not random graphs, they are locally structured, and this structure can be exploited for contact prediction. As was pointed out in [17], the density of contacts is higher in the neighbourhood of a contact than in the neighbourhood of a non-contact.

Our questions start from this observation: can we use local contact patterns to construct simple and interpretable supervised approaches to contact prediction? Can we use such approaches in the case of inter-protein and inter-domain contacts, which are less represented in the protein data bank PDB [18] than intra-domain contacts and thus provide less training data?

To this end, we introduce FilterDCA: we first analyse contact patterns related to different secondary structure elements, and show that these are faithfully reflected by the scores derived using plmDCA (i.e. DCA based on pseudo-likelihood maximisation) [19]. We therefore use average contact patterns as explicit filters for the DCA predictions, and combine them with the standard DCA score using simple logistic regression. Surprisingly, this fully transparent scheme reaches perfomances comparable to the CNN based contact predictor PConsC4 [20]. While our learning and testing procedure uses domain-domain interactions inside multi-domain proteins, due to their better availability in the PDB, we show that the prediction of inter-protein contacts is significantly improved, too.

We think that our results are interesting, because they allow to take a step forward in understanding how contact information is actually hidden in sequence alignments, and thus for understanding the performance reached by CNN-based contact prediction, or for refining their architecture to better take this information into account.

## Results and Discussion

To improve DCA-based contact prediction between interacting domains, we have collected an extensive dataset of more than 2500 intra-protein domain-domain interactions of experimentally known protein structures, containing more than 6 10^5^ inter-domain contacts, cf. *Materials and Methods*. For each of these domain pairs, a multiple-sequence alignment (MSA) is created, with each line containing the aligned sequences of both interacting domains. According to the effective number *M*_eff_ of sequences (as calculated by plmDCA in standard settings), we distinguish three cases: large (*M*_eff_ *>* 200), medium (50 *< M*_eff_ *<* 200), and small (*M*_eff_ *<* 50) MSAs. These data will be used throughout the paper, if not explicitly stated differently.

### DCA scores reflect typical inter-domain contact patterns

While DCA uses rather sophisticated approaches for global statistical modeling of MSAs of families of homologous proteins, the subsequent residue-residue contact prediction is surprisingly naïve: coevolutionary coupling matrices *J*_*ij*_(*a, b*) of dimension 21 × 21 between two residue positions *i* and *j* (MSA columns) are compressed into a single ad-hoc score, either using direct information *DI_ij_* [1] or, more frequently, by calculating a statistically corrected Frobenius norm *F*_*ij*_ [19], and the largest scores are interpreted as potential contacts. As is shown in *Materials and Methods*, Fig. 7, this strategy is informative about contacts in the case of large MSAs, becomes questionable in the case of intermediate MSAs, and totally loses signal for small MSAs.

This strategy might anyway be optimal if contact maps were totally random. However, contact maps show important patterns, mainly based on secondary structures: *β*-sheets are characterised by diagonal patterns, *α*-helices give rise to almost periodic patterns with a period of 3-4 corresponding to one helical turn: when a pair (*i, j*) is in contact, the residues *i*±2 (resp. *j*±2) are on the opposite face of the helix, and thus probably do not form contacts with any residue close to *j* (resp. *i*), while *i*±3, 4 (resp. *j*±3, 4) are again on the same face as *i* (resp. *j*).

To exploit such patterns, we have classified all residues according to their secondary structure into three classes: helical (H), extended (E) and other (O). Each inter-domain contact can now be classified by the secondary-structure annotation; in the following we will concentrate exclusively on the classes HH of two helical residues in contact, and EE for two contacting residues which are both located in *β*-strands. All other classes do not show very interesting contact patterns; their inclusion in our procedure does not lead to an improved contact prediction.

For both groups, HH and EE contacts, we calculate average contact maps over windows centered in all contacts (*i, j*), which are realised in the above-mentioned inter-domain interaction dataset for large families, cf. *Materials and Methods*. For a window size of 15 × 15, we show the results in Fig. 1.A; the entries give the estimated probabilities to find another contact in the corresponding position relative to the given contact (*i, j*) in the center of the window. While the case of HH contacts shows a clear pattern with the expected periodicity of 3-4 positions, i.e. in coherence with one turn of the *α*-helices, the case of EE contacts shows a non-informative pattern, where the contact density decays simply with the distance from the central EE contact.

**Fig 1.**
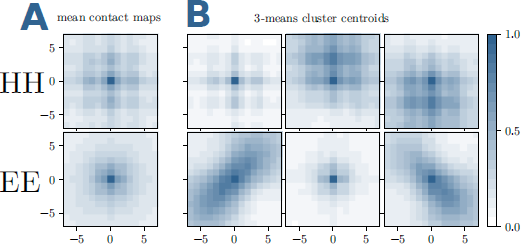
Typical patterns in inter-domain contact matrices. (A) The frequencies of contacts in a 15 × 15 contact-map window around an HH or EE contact is displayed. The average is done over the 46978 HH and 12281 EE contacts of the training set. The mean contact matrices are a combination of parallel, anti-parallel or mixed HH and EE contacts. (B) We disentangle them with a *k*-means clustering with *k* = 3. The 6 resulting centroids, 3 for a central HH inter-domain contact (upper row) and 3 for an EE contact (lower row), show pronounced patterns and can be used to filter DCA predictions.

The picture changes when we refine the analysis using *k*-means clustering with *k* = 3 of the HH and EE contact matrices. The HH patterns remain similar, unveiling some fine structure in the HH contact ensemble. The EE case now becomes highly informative, we clearly observe the two diagonal patterns corresponding to parallel resp. anti-parallel *β*-strands. The third cluster assembles all other cases, like crossing *β*-strands.

Are these contact patterns reflected by the matrices of DCA scores? To answer this question, we calculate average DCA-scores for windows centered around exactly the same contacts as those used in Fig. 1. Doing so, we average out site specificities and noise, and only coherent local patterns remain visible. As becomes evident in Fig. 2, the resulting patterns have the same structure as the averaged contact maps. However, it becomes also evident that the average DCA scores are very small, as compared to DCA scores, which are indicative for contacts according to Fig. 7.

**Fig 2.**
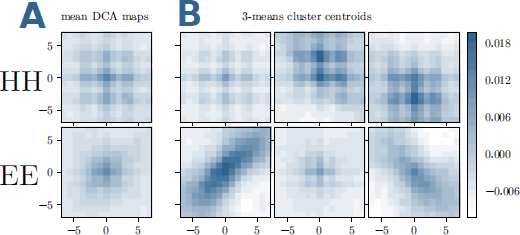
Typical patterns in DCA-score matrices. (A,B) The average DCA scores in a 15 × 15 window around an HH or EE contact are displayed, using the same selection of contacts as in Fig. 1 for panels A, and the same sub-clustering for panels B.

This double observation is the starting point for our supervised contact predictor in FilterDCA: while many contacts have small DCA scores, i.e. they would not be predicted as contacts in standard plmDCA, they can still have a characteristic DCA pattern in their neighbourhood, and thus potentially be identified as contacts.

### Integrated contact patterns and DCA improve inter-domain contact prediction

To this aim, we have developed a simple and transparent strategy to integrate plmDCA with the structural filters to improve inter-domain contact prediction, cf. Fig. 3. The idea is to integrate the standard plmDCA score, which is known to be a good contact predictor when assuming large enough values, with a coherence measure between our contact filters and the corresponding window of plmDCA scores.

**Fig 3.**
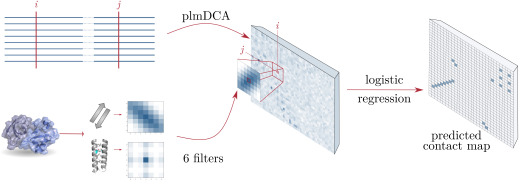
General scheme of FilterDCA: Our approach combines the results of plmDCA applied to two-domain MSAs with structural filters constructed as average contact matrices using 6 contact classes. Structural supervision is used to learn a logistic regression based on the plmDCA score itself, and the best correlation with one of the six structural filters.

To this end, we define two informative features **x** = (*x*_1_*, x*_2_) for each pair of residues (*i, j*), as a function of the DCA scores and the structural filters

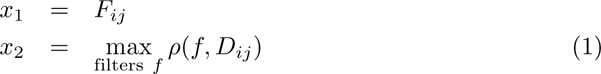

with *D*_*ij*_ being a window of DCA scores around residue pair (*i, j*), and *f* denoting the 6 filters (which, to exclude overlaps between training and test data, are now determined using the dataset of small MSAs, cf. *Material and Methods*), and *ρ* being the Pearson correlation. Note that in both *D*_*ij*_ and *f* the central element has been removed, since it would introduce a strong redundancy with the first feature. The size of the window is a free parameter whose influence will be examined. In the following, we will refer to *x*_1_ as “DCA score”, and to *x*_2_ as “filter score”.

To integrate these two features into a single contact predictor, we use simple logistic regression. The probabilities to belong to the class *contact* (⊕) or *non-contact* (⊖) are thus given by:

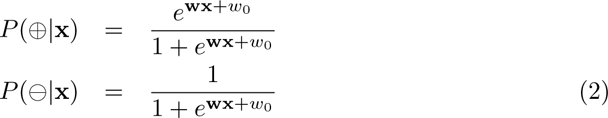

where the bias *w*_0_ and the weights **w** = (*w*_1_*, w*_2_) are optimised using a training set of 50% of the data. This is done independently for the large and intermediate MSAs, cf. *Materials and Methods* for the details of the implementation.

A first insight into the relative importance of the two features – the standard DCA score and the local coherence with the typical contact patterns measured by the filter score – can be gained from Fig. 4. For small filter sizes, the decision boundary is almost horizontal, i.e. the decision is almost exclusively determined by the DCA score, and the filter score has little influence. The decision boundary is located mainly between 0.2 and 0.3, in perfect agreement with the crossing points of the histograms of contacts and non-contacts in Fig. 7.

**Fig 4.**
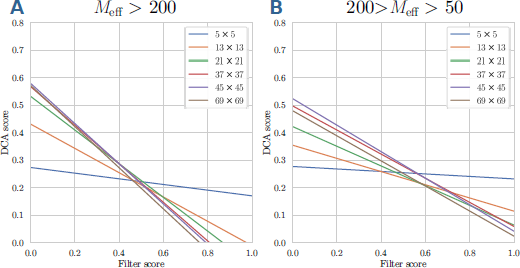
Decision boundary for logistic regression: The lines show, for large (panel A) and medium (panel B) MSA sizes, the decision boundary defined by *P* (*⊕|***x**) = 1*/*2, for different filter sizes raging from 5 × 5 to 69 × 69.

This changes when larger windows are used as filters. The decision boundary becomes tilted. Not only pairs (*i, j*) of smaller DCA score can be predicted to be contacts when being in an environment of high filter score, but also relatively large DCA scores may be discarded when not being related to a reasonably large filter score. Coherence of the DCA signal around a residue pair with local contact patterns thus has the potential to discover otherwise discarded contacts, and to prune large DCA scores judged to be isolated noise due to an incoherent environment.

Is this potential actually realised and leads to better contact predictions in the test set of protein families, which were not used for model learning? Fig. 5 shows the positive predictive value (PPV = TP/(TP+FP)) as a function of the number of predictions (TP+FP), average over all families in the test sets for large and intermediate MSA sizes. A comparison with standard plmDCA shows that the application of structural filters effectively improves the inter-domain contact prediction. However, the quality of the first predicted contacts does not change a lot, i.e. it is still dominated by the quality of the plmDCA prediction. On the contrary, the decay of the PPV with the number of predicted contacts is substantially slower. The maximal PPV is reached for quite large filters (size 45 × 45), but then it decreases again, as testified by the curve for 69 × 69 filters. Intuitively we thus find that FilterDCA is able to help only if DCA alone finds some contact signal. Coherence with contact patterns over quite large environments of the considered residue pairs (*i, j*) is most informative, but even larger filters lead to a decay since the filters take into account too distant and thus too structurally-variable regions.

**Fig 5.**
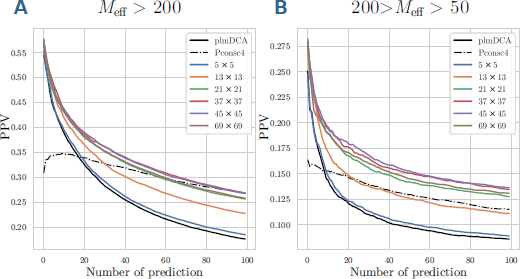
Positive predictive values of FilterDCA for inter-domain contact prediction. PPV are shown as a function of the number of predictions for large (panel A) and medium (panel B) MSA sizes, averaged over the different domain-domain interactions in the test set. Different filter sizes are compared to standard plmDCA and to the deep-learning-based PconsC4.

Interestingly, the performance is comparable to the one of PconsC4, a deep-learning-based contact predictor. While initially being slightly worse due to some probably systematic mispredictions of PconsC4, the asymptotic behavior of PconsC4 is very similar to the one of FilterDCA, with a small advantage for PconsC4 in the case of large MSA, and for FilterDCA in the case of intermediate MSAs. However, on one hand it has to be noted that PconsC4 was trained on intra-domain contacts, and applied to inter-domain contacts, a fact which might systematically impact on the prediction quality of PconsC4. On the other hand, PconsC4 is trained on much more contacts than we need for FilterDCA. It would be interesting to compare FilterDCA also to other deep-learning-based contact predictors like RaptorX, but we were not sure about the precise training set of the RaptorX web server, i.e. about the independence of the training set of RaptorX from our test set. We would like to repeat, however, that FilterDCA was not designed in the first place to be the most performant contact predictor, but to be fully transparent and interpretable.

### Training on intra-protein / inter-domain contacts improves inter-protein contact prediction

In the introduction, we have motivated our work on domain-domain interactions and predicting inter-domain residue contacts by the idea that these interactions can actually serve as proxies for protein-protein interactions (PPI) and inter-protein contact prediction, but have more representatives in the structural databases, thus allowing for better structure-based supervision. Is this actually true?

To answer this question, we have used the PPI dataset of [21] as a test set, applying Filter DCA as learned on the intra-protein / inter-domain dataset used in the last section (case of large MSAs), for technical details cf. *Materials and Methods*. The results are depicted in Fig. 6. We find that FilterDCA can be robustly transfered from inter-domain to inter-protein contacts, with comparable gains in PPV in the two cases (the general performance seems a bit higher, but we guess it is a direct effect of the smaller and curated PPI dataset in [21]). Interestingly, PconsC4 seems to suffer more from the transfer to PPI, the initially weak performance being even more pronounced than in the inter-domain case. This illustrates that the transfer itself is not trivial, with inter-domain contacts representing an intermediate case between intra-domain and inter-protein contacts.

**Fig 6.**
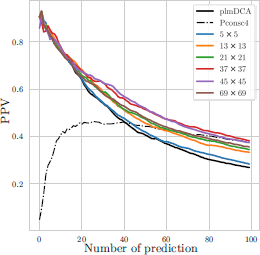
Positive predictive values of FilterDCA for inter-protein contact prediction. PPVs are shown as a function of the number of predictions, averaged over the different protein-protein interactions in the test set. Different filter sizes are compared to standard plmDCA and to the deep-learning-based PconsC4.

## Conclusion

Despite some success, the direct applicability of DCA and related coevolutionary methods in predicting contact maps and thereby allowing to model tertiary and quaternary protein structure has remained limited. The reason is the fully unsupervised character of DCA – only sequence information is used, and the relation between sequence-related coevolutionary coupling scores and protein structure was based on the simplistic assumption, that large individual scores predict contacts. However, most contacts show quite weak coevolutionary signals, which are comparable to the coevolutionary couplings between non-contacting residues, which are inferred to be non-zero, e.g., because of the finite size of the training MSA and phylogenetic correlations between the sequences in the MSA.

Structure-based supervision had been applied with impressive success to overcome this limitation. In particular deep-learning, mostly using convolutional neural networks, has been able to establish much more accurate relations between coevolutionary couplings and residue contacts, in particular by integrating local sequence features like predicted secondary structure and solvent accessibility. The complexity of deep neural networks makes, however, the interpretability of the results very hard, and the high number of parameters in deep neural networks makes large training sets essential.

In this work, we have taken a complementary approach to introduce structure-based supervision. Instead of relying on the capacity of flexible machine-learning tools to learn even complex patterns from sufficiently abundant data, we have benefitted from our prior biological knowledge to explicitly construct features, which are informative about contacts beyond individually large DCA scores. To this aim, we have used typical contact patterns as explicit filters for the environment of each residue pair – we have found that the strongly non-random structure of contact maps provides valuable contact information if used to filter the DCA predictions.

This approach has the advantage of being robust even for limited datasets (we have used domain-domain interactions), but it has itself obvious limitations: building our supervision on explicitly constructed contact patterns, we cannot find alternative, unexpected and thus surprising informative features. More flexible architectures like convolutional neural networks may do so, but at the expense of many additional parameters to be learned from the training data.

It would now be interesting to see how far other biological features are informative about residue-residue contacts. At least three possibilities may come directly to our mind: first, surface exposure may be a very interesting feature, in particular for protein-protein interactions between compact domains – only residues exposed at the surface of the monomers have a reasonable chance to form inter-protein contacts. Second, not all types of amino acids are biophysically compatible to form stabilising contacts. DCA couplings reflect such biophysical interactions, but a direct implementation of amino-acid interaction matrices might contribute to an improved contact prediction. Third, interaction interfaces form typically a limited number of connected patches on the protein surfaces – this may be used as a coherence measure between different predicted contacts. Each of these may give a contribution to contact prediction, leaving room for the future exploration of interpretable coevolution-based contact predictors.

## Materials and methods

### Domain-domain interaction dataset

The basis of our dataset is given by the 3did database [22], which was constructed by selecting high-resolution PDB structures [18] containing multiple contacting Pfam domains [23]. We use the Aug 5, 2017 version which is based on Pfam v.30.0 and contains a total of 11.200 structurally-resolved domain-domain interactions. To get the joint MSAs we exclude homodimeric cases, and match sequences of domains co-localized on the same protein chain, i.e. we consider exclusively intra-protein inter-domain interactions. Finally, we map each residue in the MSAs to the corresponding positions in the PDB files. The mapping is done by aligning the PDB sequences to the profile HMMs of the PFAM domains through hmmalign [24], which allows to associate residue-residue distances to any pair of alignment columns.

Often only a part of the MSA can be mapped to the corresponding PDBs. We keep only MSA s with domain mapping coverage greater than or equal to 40% of the Pfam-domain length. The distance between two residues is defined as the minimal distance between all heavy-atoms of the two residues; in case the same residue-residue pair is associated with multiple PDBs, we assign the minimum distance between all possible copies. This assumes that any predicted contact (distance below 8Å), which is present in at least one PDB structure, is a true positive prediction. We further clean our dataset by requiring at least 10 and at most 2000 residue-residue interactions and removing few cases of coiled-coil structures which, due to repeated motifs, can lead to spurious coevolutionary signal.

At the end, we keep a total 2598 joint MSAs of pairs of contacting domains. A list of these domain-domain interactions is provided in the Supplementary data.

### Direct coupling analysis

We have run plmDCA [19] in A. Pagnani’s Julia implementation (available at https://github.com/pagnani/PlmDCA) with standard settings. Two outputs are of relevance: for each pair (*i, j*) of residues a DCA score *F*_*ij*_ is provided (via the statistically corrected Frobenius norm of the DCA coupling matrices), and the effective sequence number *M*_eff_. With the scope of predicting inter-domain contacts, we only consider residue position pairs with *i* in the first, and *j* in the second domain.

It is well known that the accuracy of DCA predictions is strongly dependent on the number of sequences in the MSA or, more precisely, on the effective number of sequences *M*_eff_ - the higher the better. DCA predictions are comparable only for MSAs having similar *M*_eff_. Thereby, we decide to split the 2598 MSAs of contacting domains in 3 datasets according to *M*_eff_, see Table 1, and to analyse them independently. As can be seen in Fig. 7, which shows histograms of DCA scores for all contacting and non-contacting residue pairs, only for the largest *M*_eff_ a clear contact signal is contained in the largest DCA scores.

**Fig 7.**
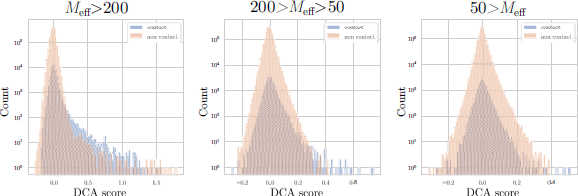
The distributions of DCA scores of contacts and non-contacts. For MSAs with *M*_eff_ > 200 (Panel A) and 50 *< M*_eff_ ≤ 200 (Panel B). Note that the enrichment of true positive predictions (contacts) is very high in the tail of large DCA scores. In fact, the majority of the pairs with DCA score larger than 0.3 corresponds to contacts. This is not the case for MSA with *M*_eff_ *<* 50 (Panel C), where the two distributions completely overlap.

**Table 1.**
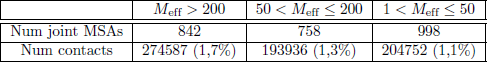
We split the 2598 MSAs of interacting domains in 3 datasets according to the effective number of independent sequences, *M*_eff_. They contain approximately the same number of MSAs (first row) and inter-domain contacts (second row). In brackets, the percentage of inter-domain contacts is given with respect to the total number of inter-domain residue pairs.

| | $M_{\text{eff}} > 200$ | $50 < M_{\text{eff}} \leq 200$ | $1 < M_{\text{eff}} \leq 50$ |
| --- | --- | --- | --- |
| Num joint MSAs | 842 | 758 | 998 |
| Num contacts | 274587 (1,7%) | 193936 (1,3%) | 204752 (1,1%) |

### Filter score

With the aim of going beyond simple DCA predictions, we define a new score by applying structural filters on DCA predictions, cf. Fig. 3. Let

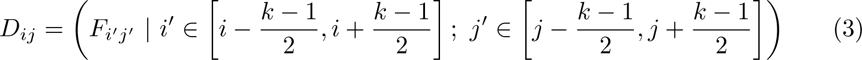

be the matrices of size *k × k* of inter-domain DCA scores, centered in residue pair (*i, j*) (i.e. windows of the full plmDCA output). We always choose *k* to be an odd number in order to get a square matrix centered around the central pair, in practice we use window sizes *k* between 5 and 69.

The construction of the structural filters *f ∈ **S*** is as described in the main text: HH and EE inter-domain contacts are clustered into three clusters each, by using 3-means clustering. The filters are the centroids of the corresponding clusters. For all learning tasks (large and medium MSAs), we have used the same filters, determined using the contacts in the dataset of small MSAs, cf. Fig. 8 for *k* = 13. Note that the filters are very similar to the ones in Fig. 1, which were determined using all contacts in the large-MSA dataset. This illustrates the robustness of the filter construction.

**Fig 8.**
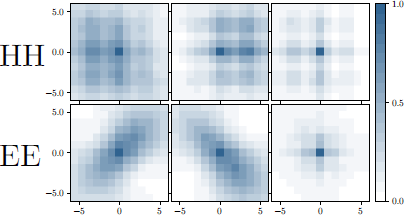
Filters used in the calculation of the filter score. The filters were determined from the inter-domain contact maps in the small MSA, with *k*-means clustering for *k* = 3 applied separately for HH and EE contacts.

For each of the 6 filters *f ∈ S* of size *k × k*, we compute the Pearson correlation between each *D*_*ij*_ and the filter. The central pair is removed from the calculation since, for the filters in ***S***, it is a contact by construction and since the DCA score of the central pair (*i, j*) will be used directly. The new score, named *filter score*, equals the maximum of the 6 Pearson correlations.

A problem arises from pairs of residues closer than (*k −* 1)*/*2 to the border of the DCA matrix: the matrix *D*_*ij*_ is smaller than the filter matrix, and the procedure displayed in Fig. 3 cannot be directly applied. In this case we compute the Pearson correlations only for pairs which are both contained in *D*_*ij*_ and in the filter matrix.

### Learning procedure

FilterDCA is based on supervised logistic regression which takes as input two features: (i) the standard DCA score, and (ii) the previously defined filter score, cf. Eq. (1). It outputs the probability for a pair of residues to be in contact, as given in Eq. (2). The bias *w*_0_ and the weights **w** = (*w*_1_*, w*_2_) are optimised using the ‘liblinear’ solver of the *sklearn* library [25].

Pairs of residues forming a contact are only a small fraction of all possible pairs, cf. Table 1). Thus, the training set is strongly imbalanced: the incidence of the *non-contact* class is dominant, being found in 99% of cases. We found that the performance of our classifier is improved if we restrict the training set to residue pairs with DCA score *F*_*ij*_ larger than zero, cf. Fig. 7. In this case, the classifier concentrates on cases which show a more reliable coevolutionary signal, and which are concentrated closer to the decision boundary. Another further slight improvement has been achieved by scaling the filter scores, in both training and test set:

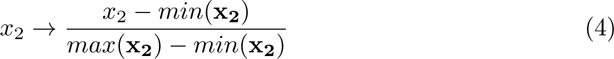

where *max*(**x_2_**) [*min*(**x_2_**)] is the maximum [minimum] filter score in the training set. While the classifier itself is invariant under this transformation, the *ℓ*_2_-regularisation used by *sklearn* is not, thereby influencing the final parameter values.

## Acknowledgments

We thank Juliana Silva Bernardes for interesting discussions during this work. MW acknowledges funding by the EU H2020 research and innovation programme MSCA-RISE-2016 under grant agreement No. 734439 INFERNET. This work has been undertaken partially in the framework of CALSIMLAB and supported by the public grant ANR-11-LABX-0037-01 overseen by the French National Research Agency (ANR) as part of the “Investissements d’Avenir” programme (ANR-11-IDEX-0004-02).

